# Pupillometry reveals autonomic adjustments during diving reflex in face immersion apnea

**DOI:** 10.1101/2025.01.31.635967

**Authors:** V. Rizzuto, R. Montanari, L. Mesin, M. H. Bortolozzo-Gleich, M. Laurino, Y. Bonneh, D. Yellin, G. Laganovska, J. Vanags, G. Giuliano, E. Moccia, M. Zanoletti, P. Bufano, E. Melissa, MV Uthaug, V. Lionetti, P. Longobardi, A Gemignani

**Affiliations:** School of Advanced Science, University of Camerino, Camerino, Italy; Institute of Pharmacology, Heidelberg University Hospital, Heidelberg, Germany; Mathematical Biology and Physiology, Department of Electronics and Telecommunications, Politecnico di Torino, Turin, Italy; Instituto de Neurociencias UMH-CSIC (Alicante), Avenida Santiago Ramón y Cajal s.n., 03550, Spain; Institute of Clinical Physiology, National Research Council, Pisa, Italy; School of Optometry and Vision Science, Faculty of Life Science, Bar Ilan University, Ramat Gan, Israel; Department of Brain Sciences, Weizmann Institute of Science, Rehovot, Israel; Clinic of Ophthalmology, P. Stradins Clinical University Hospital, Riga, Latvia; Department of Ophthalmology, Riga Stradins University, Riga, Latvia; Pegaso University, Faculty for Human Sciences, Rome, Italy; Latvian American Eye Center (LAAC), Riga, Latvia; Master II level Underwater and Hyperbaric Medicine “Piergiorgio Data”, Scuola Superiore Sant’Anna, Pisa, Italy; Department of Surgical, Medical and Molecular Pathology and Critical Care Medicine, Pisa, Italy; Department of Neuroscience, University of Pisa Hospital, Pisa, Italy; Department of Neuropsychology and Psychopharmacology, Faculty of Psychology and Neuroscience, Maastricht University, Maastricht, Netherlands; Centre for Psychedelic Research, Department of Brain Sciences, Imperial College London, London, United Kingdom; Unit of Translational Critical Care Medicine, Laboratory of Basic and Applied Medical Sciences, Interdisciplinary Research Center “Health Science,” Scuola Superiore Sant’Anna, Pisa, Italy; Hyperbaric Centre, 48124 Ravenna, Italy

**Keywords:** diving reflex, pupillometry, hippus

## Abstract

The human diving reflex is an innate cardiorespiratory adjustment triggered during apnea that concurrently activates both sympathetic and parasympathetic branches of the autonomic nervous system to maintain physiological stability under stress. Although pupil dilation and constriction are antagonistically regulated by these branches, the effect of the diving reflex on pupil diameter oscillations, known as hippus, remains unexplored. Here, we compared hippus in healthy participants during breathing or apnea either with (Wet) or without (Dry) facial immersion in cold water. In both apneic conditions, hippus exhibited reduced power in the low-frequency band (< 0.25 Hz). Notably, during Wet apnea, we observed a reallocation of power towards higher frequencies (> 0.25 Hz) and increased entropy of fluctuations, indicating a shift in autonomic balance and greater signal complexity during the diving reflex. This pilot study highlights pupil dynamics as a sensitive and non-invasive probe of autonomic adjustments underlying the human diving reflex.

## Introduction

Pupil diameter is regulated by the coordinate activity of the parasympathetic (PNS) and sympathetic (SNS) branches of the autonomic nervous system^[1]^. Through its innervation of the iris sphincter muscle, the PNS induces pupil constriction, whereas the SNS promotes dilation via activation of the radially oriented dilator muscle^[1]^. Although these two pathways are largely independent, their output can be centrally coordinated by the locus coeruleus ^[2]^. Acting on the Edinger-Westphal nucleus, the locus coeruleus can indirectly inhibit PNS-mediated constriction while directly facilitating the SNS-driven dilation. The continuous interplay between the two branches produces spontaneous oscillations in pupil diameter, known as pupil unrest or hippus^[3]^. Hippus typically fluctuates within a frequency range of 0.047 to 2 Hz ^[4]^, with a dominant peak around 0.3 Hz ^[3]^, leading to diameter changes up to over 0.5 mm. Because hippus reflects moment-to-moment adjustments in autonomic balance, it represents a promising non-invasive marker of central autonomic regulation.

The diving reflex (DR) is an evolutionarily conserved autonomic response, present across vertebrates and mammals, that enables organisms to tolerate apnea and cold-water immersion ^[5]^. In humans, the DR elicits a coordinated set of physiological reactions, including bradycardia and peripheral vasoconstriction, that involve the simultaneous activation of both PNS and SNS ^[6]^. This dual autonomic recruitment preserves oxygen supply to vital organs, particularly the heart and brain, during diving ^[5,6]^. The reflex is initiated by voluntary apnea and is further potentiated by facial cold-receptor stimulation^[7]^. Given that the DR engages both autonomic branches that control pupil size, we hypothesized that it could also modulate hippus dynamics.

To test this, we analysed pupil fluctuations in a pilot cohort of five free-divers under three experimental conditions designed to elicit progressive DR activation: 1) controlled breathing (no DR activation), 2) breath-hold without immersion (i.e., Dry apnea; weak DR), and 3) breath-hold with face immersion in cold water (i.e., Wet apnea; strong DR). We observed that Wet apnea was associated with an enhanced PNS contribution and a reduced SNS influence on hippus, together with increased signal complexity. These preliminary findings suggest that pupillometry may serve as a sensitive, non-invasive tool for tracking autonomic adjustments underlying the human diving reflex.

## Methods

### Participants

The study enrolled 5 healthy male volunteers (Supplementary Tab.1) aged between 30 and 63 years, all of whom were emmetropic and had no comorbidities or medication intake. Inclusion criteria were: non-smokers, light caffeine and alcohol consumers, drug-free for at least 2 months with no history of cardiovascular, pulmonary, or neurological/psychiatric diseases. Participants reported no history of ocular surgery, trauma, or infection, and no prior uveitis, and provided informed consent for their participation in the experiment. We included freedivers who had more than three years of freediving experience, could sustain static apnea for over three minutes, and had reached depths exceeding 25 meters.

### Experimental conditions

Breathing condition involved controlled breathing, consisting of 4 seconds of inspiration followed by 6 seconds of expiration (i.e., 6 respiratory cycles per minute). Dry involved maximal breath-holding. Wet involved maximal breath-holding too except that the face was submerged in a glass basin of water at 13 °C, constantly monitored through a mercury thermometer (Supplementary Fig.1). In all conditions, pupillometry was performed with the Braineye eye-tracker (https://braineye.com/science/), shining an IR light (850 nm), plus a BioEye NIR camera built to optimise pupil acquisition by preserving wavelengths that enhance pupil-to-iris contrast. Pupil diameter values were extracted at ∼30 Hz and streamed to a Xiaomi 9 mobile. Heart rate (BPM) and peripheral oxygen saturation data were collected using Gima oxy-2 and Gima oxy-3 pulse oximeters at a sampling rate of Hz. Free-divers were asked to stare at the fixation point located at 40 cm distance on the surface of the eye-tracker, while lying ventrally in a customised operating table resting the forehead and chin on a headrest assembly (Supplementary Fig.1). The illumination of the room was set at 70 ± 5 lumen.

### Data curation and analysis

BioEye software provides a median pupil score (MPS) to rate the recording quality. This metric is defined by the sharpness of the iris-to-pupil contrast in the obtained image, where a threshold value >= 8 indicates an excellent separation between the two. Therefore, recordings were excluded if MPS <8. Additionally, from the two eyes, we only selected the recording of the eye with the lower standard deviation in iris size variations. Recordings were analysed with Matlab 2023a. Given the non-uniform sampling of the ∼30 Hz signals, the recordings were resampled to 30 Hz using piecewise cubic interpolation. Blinks were removed from the pupil diameter signal, and the resulting trace was interpolated (again using a piecewise cubic interpolation) to construct a continuous signal (joining the otherwise separated segments of epochs without blinks). Subsequently, data were band-pass filtered between 0.04 and 2 Hz (Chebyshev type II filter, with 0.5 dB of maximum ripple and 25 dB of minimum attenuation below 0.02 Hz and above 2.5 Hz). The power spectral density (PSD) was estimated using Welch’s method, applied to 10-second temporal windows (Tukey window with cosine fraction of 0.1) with 50 percent overlap. Then, the power associated with the signal for each band was obtained by integration. Approximate Entropy (ApEn), an estimation of time-series complexity ^[8]^, was computed with a small embedding dimension (i.e., 2), after removing low-frequency trends and resampling the signals at about 3 times their bandwidths. Specifically, the low-frequency component was obtained using a Chebyshev type II low-pass filter with 0.5 dB maximum ripple and 20 dB minimum attenuation, with a transition band between 0.24 and 0.27 Hz; the high-frequency component was defined as the difference between the signal and the low-frequency component. Then, the two components were resampled at 0.8 and 8 Hz, for the low and high frequencies, respectively. To reduce the effect of self-recurrences in the signal ^[9]^, we calculated a modified version of the ApEn, in which the tolerance value was chosen to select a number of recurrences equal to a percentage of the total number of samples (10% in this work).

### Statistics

To compare the power spectral density and approximate entropy percentages across the 3 experimental conditions, a one-way analysis of variance (ANOVA) was used after testing for normality (Shapiro-Wilk test). As we performed multiple observations on the same subjects, we used nested statistical testing to account for the reduced degrees of freedom. In the case of statistical significance, this was followed by a multiple-comparison test (Tukey-Kramer test), providing adjusted p-values for pairwise comparisons. The alpha level was set at 0.05.

## Results

Free-divers rested on a custom-made operating table, lying ventrally, with the chin and the forehead supported by a headrest assembly. In Wet, they would additionally submerge the face in the cold water of a basin beneath it (Supplementary Fig. 1). DR occurrence was monitored by simultaneously measuring blood oxygen saturation (SaO_2_) and heart rate. They achieved an average Dry apnea duration of 139 ± 15.13 seconds and an average Wet apnea duration of 150.8 ± 29.86 seconds. Despite the low levels of SaO_2_ they reached at the end of apnea (87,53 % ±10,9 % in Dry, 93,03 % ± 6,73 % in Wet) no subject had any medical and/or neurological complication as a consequence of the apnea. Consistent with other descriptions ^[33, 46, 47]^, heart rate showed a decrease-increase pattern [mean ± S.E.M. (BPM): start: 66.9 ± 1.5, middle: 62.3 ± 1.8, end: 68.2 ± 2.3] and pupil diameter showed the typical hippus (Fig. 1a,b). Across single performances, pupil diameter (Fig. 1c) tended to be significantly smaller in Wet than in Dry and Breathing conditions (absolute medians’ difference (mm): Breathing - Dry= 0.08; Breathing - Wet= 0.54; Dry - Wet= 0.62) [One-way ANOVA: F(2,27)= 4.03, p= 0.0293], however post-hoc analysis only indicated tendencies for pairwise differences [p_Breathing *vs* Dry_= 0.9576, p_Breathing *vs* Wet_= 0.0720, p_Dry *vs* Wet_= 0.0634], possibly due to low sample size. Also, Power Spectral Density (PSD) of hippus in both Dry and Wet showed significantly lower power in the low-frequency band than Breathing (Fig 1d,e) [Nested one-way ANOVA: F(2,27)= 8.95, p= 0.001; Tukey’s multiple comparison: p_Breathing *vs* Dry_= 0.0054, p_Breathing *vs* Wet_= 0.0009, p_Dry *vs* Wet_= 0.8272], while net high-frequency power did not differ (Fig 1d,f) [Nested one-way ANOVA: F(2,27)= 2.25, p= 0.147]. Yet, the %Power, defined as the fraction of power for low- and high-frequencies over the total power within the 2Hz range, was significantly higher in the low than the high-frequency band for Breathing [Nested t-test: F(1,10)= 77.46, p< 0.0001] and Dry [Nested t-test: F(1,8)= 11.39, p = 0.0097]. Instead, Wet exhibited the opposite pattern, with the high-frequency band containing significantly %Power compared to the low-frequency band [Nested t-test: F(1,8)= 15.09, p = 0.0046], suggesting the occurrence of a power re-allocation in this condition. Lastly, we explored the hypothesis that the DR could affect the complexity of the hippus. Indeed, we found that the Approximate Entropy in Wet was significantly higher than Breathing and Dry (Fig. 1h) [Nested one-way ANOVA: F(2, 27) = 19.89, p <0.0001; Tukey’s multiple comparison: p_Breathing vs Dry_= 0.196, p_Breathing vs Wet_ <0.0001, p_Dry vs Wet_= 0.0003].

**Fig 1.**
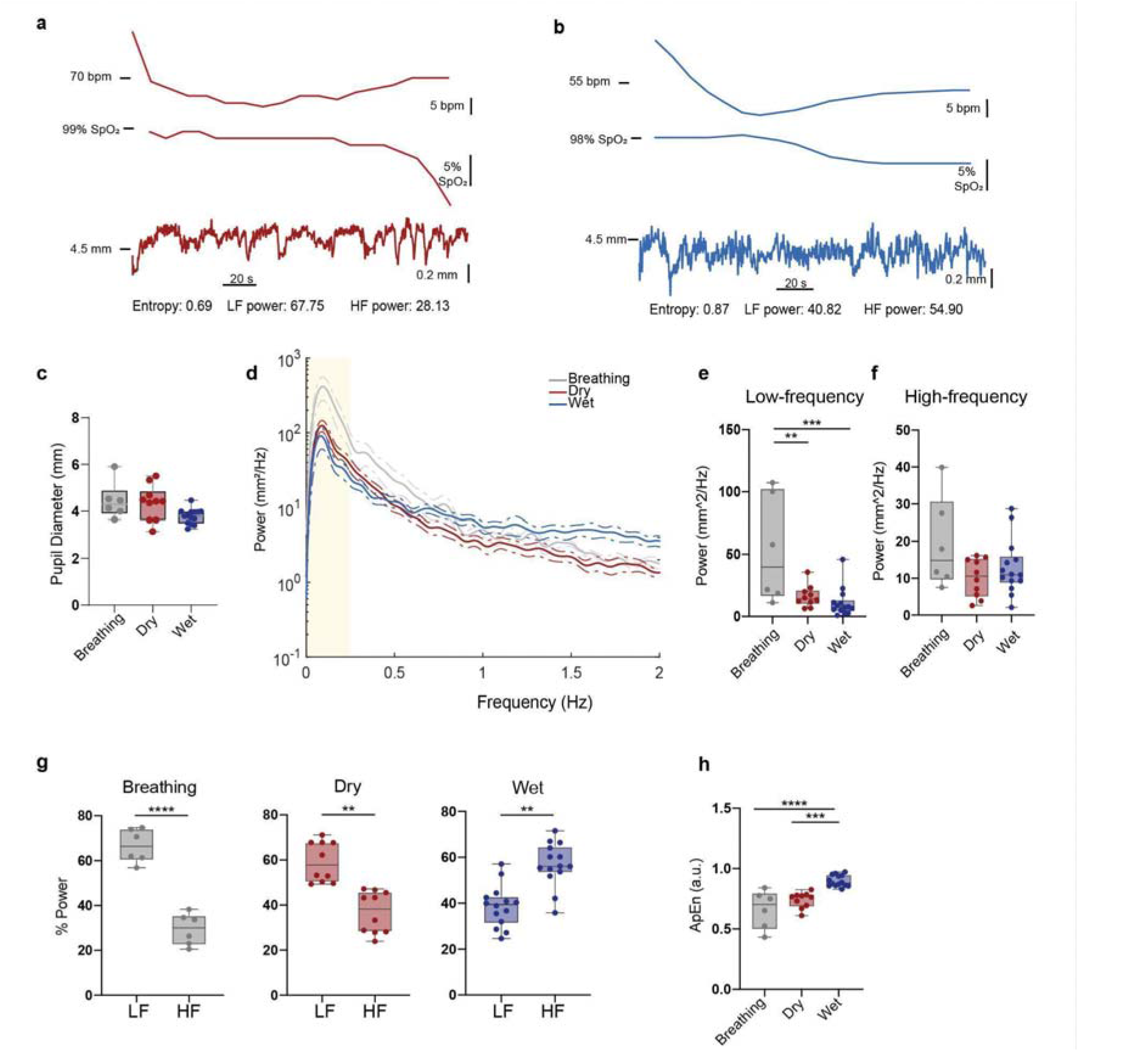
Dry and Wet apnea modulate the hippus. a) Example parallel recording of heart rate (upper panel) and oxygen saturation (middle panel) associated with pupil diameter fluctuations and its characterisation (bottom panel) during apnea in Dry condition. b) Same as *a* for apnea in the Wet condition. c) Average pupil diameter in the three experimental conditions. d) Power spectral density (PSD) semi-log plot of pupil diameter fluctuations in the three experimental conditions ± S.E.M (broken lines). Shaded area indicates the selected sympathetic frequency band. e) Statistical comparison of Power contained in the low-frequency band in the three conditions. Each data point represents a single performance. f) Same as *e* for the high-frequency band. g) Comparison of %Power between low- and high-frequencies for each experimental condition. h) Comparison of Approximate Entropy in the three experimental conditions. LF= low-frequency; HF= high-frequency.

Thus, by revealing changes in hippus dynamics across our experimental conditions, these results suggest that pupillometry is a sensitive metric for studying the DR.

## Discussion

The attempt to associate specific frequency components of hippus with distinct autonomic processes, including respiration, cardiac rhythm, and emotional arousal, has led to various classification schemes linking different frequency bands to PNS and SNS influences ^[3,10,11,12]^

This diversity reflects both methodological heterogeneity and the complexity of central autonomic integration, in which PNS and SNS activities often overlap rather than alternate strictly. In the present study, we adopted a frequency-based classification grounded in pharmacological evidence. Specifically, we attributed the 0.04-0.25 Hz band to SNS activity and the 0.25-2 Hz band to PNS modulation, based on the findings of Turnbull et al. ^[3]^, who reported a marked reduction of energy within the 0.24-1 Hz range after administration of tropicamide, a muscarinic antagonist that selectively inhibits PNS-mediated pupil constriction. This result provided a physiological rationale for considering about 0.25Hz as a lower boundary for PNS-driven fluctuations. Because subsequent studies have shown significant PNS effects extending beyond 1Hz ^[3, 11, 14]^, we expanded the PNS band to 0.25-2 Hz, assigning lower frequencies (<0.25 Hz) to SNS contributions. This operational separation, though simplified, allowed us to extract meaningful indices of relative autonomic balance from the spectral structure of hippus.

Our concurrent measures of heart rate during apnea, the profile of SaO_2_ at the performances’ end, and apnea duration across experimental conditions confirm a graded activation of the DR. Compared with controlled breathing, both apneic conditions (Dry and Wet) elicited bradycardia and a progressive decline in oxygen saturation (SaO□), with the Wet condition producing the longest apnea times. These physiological signatures align with stronger recruitment of the DR, consistent with classical literature showing that facial cold stimulation potentiates the reflex through activation of trigeminal afferents and vagal efferents. Consistent with this progressive DR activation, we detected a reduction in the sympathetic (low-frequency) contribution to hippus during both apneic conditions relative to the breathing baseline. This pattern supports a parasympathetic shift in ocular autonomic balance, mirroring the systemic bradycardia observed during apnea. Interestingly, only in the Wet condition was this sympathetic reduction accompanied by a significant relative increase in the PNS (high-frequency) component. The simultaneous increase in hippus entropy suggests that the system transitions from a more regular, SNS-dominated state to a more complex, adaptive autonomic configuration in which both branches interact dynamically. This observation aligns with previous studies linking increased physiological complexity to enhanced autonomic flexibility and resilience ^13^.

Taken together, our findings suggest that hippus frequency composition and entropy can serve as sensitive optical markers of the autonomic coactivation that characterizes the DR. The results extend the scope of pupillometry beyond cognitive and emotional research, demonstrating its value as a non-invasive tool for probing autonomic integration in physiological and environmental challenges. Because the pupil is under direct influence of the locus coeruleus–Edinger–Westphal circuitry, it likely provides a unique window into the central coupling between arousal and parasympathetic tone during states of hypoxia or breath-holding.

This pilot study naturally entails limitations. The small cohort size and the inclusion of trained free-divers limit generalizability to the broader population. Sex-related differences in autonomic control and pupillary dynamics were not assessed, and interindividual variability in training or stress response may have influenced results.

Future studies should therefore enlarge and diversify the cohort, including both sexes and different experience levels, and systematically manipulate respiratory patterns (i.e.: free breathing, voluntary hyperventilation, or slow breathing).

Integrating psychometric assessments, EEG, or heart rate variability analysis will help delineate the full autonomic and emotional imprint of the diving reflex on pupillary dynamics. Such multimodal designs could clarify whether hippus complexity reflects merely peripheral adjustments or broader reorganization of the central autonomic network.

In conclusion, our preliminary findings indicate that pupil dynamics, when analyzed in the frequency and entropy domains, encode measurable signatures of autonomic reconfiguration during the human diving reflex. Pupillometry thus emerges as a promising, non-invasive method for monitoring autonomic flexibility and adaptation in physiological, environmental, and clinical contexts.

---

**Supplementary Fig 1.**
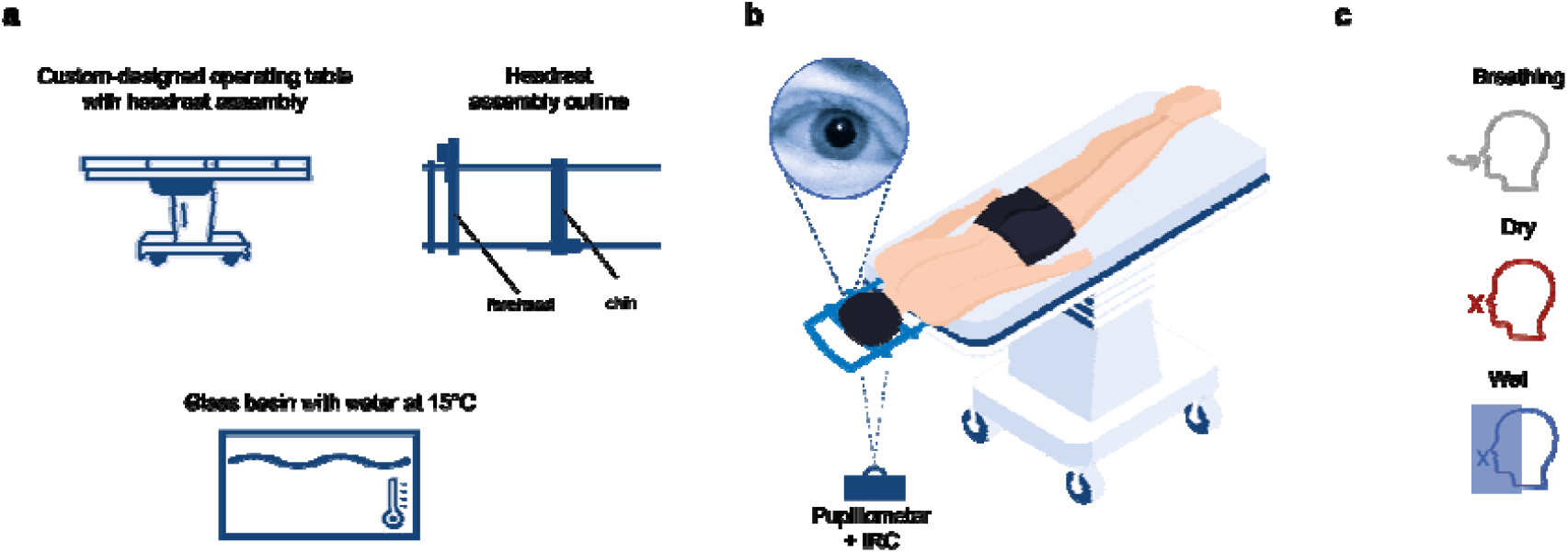
Experimental setup. a) Overview of the tools used to position the subjects. Care was taken for the subjects to feel comfortable and decrease pressure on the chest. Water temperature was constantly monitored through an internal thermometer. b) Illustration of the positioning of subjects, who were instructed to lie face down on the operating table with their heads centered and stabilized using the headrest. The eye tracker was positioned beneath the subjects’ faces, and a calibration system was employed. Inset shows an IRC capture of the eye in Wet condition through the water-filled transparent basin. c) Experimental conditions tested. The ‘X’ indicates absence of breathing, the arrow its presence. Note, in Wet a local anesthetic was applied to the eyes’ surface.

## References

1. McDougal DH, Gamlin PD. Autonomic control of the eye. Compr Physiol. (2015) 5:439–73. doi: 10.1002/cphy.c140014

2. Szabadi, Elemer. “Functional organization of the sympathetic pathways controlling the pupil: light-inhibited and light-stimulated pathways.” Frontiers in Neurology 9 (2018): 1069.

3. Turnbull, Philip RK, et al. “Origins of pupillary hippus in the autonomic nervous system.” Investigative ophthalmology & visual science 58.1 (2017): 197–203.

4. McLaren JW, Erie JC, Brubaker RF. Computerized analysis of pupillograms in studies of alertness. Invest Ophthalmol Vis Sci. 1992; 33: 671–676.

5. Michael Panneton, W. “The mammalian diving response: an enigmatic reflex to preserve life?.” Physiology 28.5 (2013): 284–297.

6. Foster, G. E., and A. W. Sheel. “The human diving response, its function, and its control.” Scandinavian journal of medicine & science in sports 15.1 (2005): 3–12.

7. Schagatay, Erika, Johan PA Andersson, and Bodil Nielsen. “Hematological response and diving response during apnea and apnea with face immersion.” European journal of applied physiology 101 (2007): 125–132.

8. SM. Pincus. Approximate entropy as a measure of system complexity. Proc. Natl. Acad. Sci. U.S.A., 88:2297–2301, 1991.

9. L. Mesin. Estimation of complexity of sampled biomedical continuous time signals using approximate entropy. Front Physiol., 9:710, 2018.

10. Schumann, Andy, et al. “Sympathetic and parasympathetic modulation of pupillary unrest.” Frontiers in Neuroscience 14 (2020): 178

11. Onorati, Francesco, et al. “Characterization of affective states by pupillary dynamics and autonomic correlates.” Frontiers in neuroengineering 6 (2013): 9.

12. Calcagnini, G., Censi, F., Lino, S., and Cerutti, S. (2000). Spontaneous fluctuations of human pupil reflect central autonomic rhythms. Methods Inf. Med. 39, 142–145

13. Lipsitz, Lewis A., and Ary L. Goldberger. “Loss of’complexity’and aging: potential applications of fractals and chaos theory to senescence.” Jama 267.13 (1992): 1806–1809.

14. Rizzuto, V., Laurino, M., Montanari, R., Gemignani, A., Figus, M., Covello, G., … & Mesin, L. (2025). Pupillary Hippus as a Biomarker: Spectral Signatures and Complexity Approaches in Autonomic and Clinical Contexts. Bioengineering, 12(12), 1376.

