## Supplementary figure for "Pupillometry reveals autonomic adjustments during diving reflex in face immersion apnea"

Table 1: Summary of Apnea Results

| Subject | Sex | Age | Exp. Level | Duration of Dry Apnea  (s) | Duration of Wet Apnea  (s) | Avg. PD during Breathing (mm) | Avg. PD during Dry Apnea  (mm) | Avg. PD during Wet Apnea  (mm) | Min SpO_2_ in Dry Apnea  (%) | Min SpO2 in Wet Apnea  (%) |
| --- | --- | --- | --- | --- | --- | --- | --- | --- | --- | --- |
| 1 | M | 63 | Advanced | 140 ± 17 | 175 ± 49 | 3.65 ± 0.29 | 3.23 ± 0.43 | 3.59 ± 0.35 | 69.33 ± 7.09 | 81.5 ± 6.92% |
| 2 | M | 40 | Advanced | 163 ± 20 | 180 | 4.13 ± 0.62 | 3.66 ± 0.51 | 3.66 ± 0.24 | 89.33 ± 3.21 | 95 ± 5.20% |
| 3 | M | 33 | Advanced | 140 ± 20 | 160 ± 20 | 4.45 ± 0.67 | 4.48 ± 0.44 | 3.58 ± 0.32 | 87 | 93.67 ± 1.53% |
| 4 | M | 30 | Beginner | 126 ± 15 | 126 ± 46 | 5.09 ± 0.69 | 5.52 ± 0.59 | 3.88 ± 0.15 | 96 ± 3.0 | 96 ± 2.83% |
| 5 | M | 30 | Beginner | 126 ± 11 | 113 ± 11 | 4.52 ± 0.31 | 4.48 ± 0.31 | 3.98 ± 0.09 | 96 ± 1.73 | 99 |

Table Legend for Abbreviations

Duration of Apnea: Represents the mean values of apnea performance for the subject under wet and dry conditions in seconds ± standard deviation.

PD (Pupil Diameter): Indicates the pupil diameter value expressed in millimeters ± standard

deviation.

Min SpO_2_ (Minimum Peripheral Oxygen Saturation): Denotes the minimum value of peripheral oxygen saturation at the end of apnea, expressed in percentage ± standard deviation.
